# High-throughput chromatin accessibility profiling at single-cell resolution

**DOI:** 10.1101/310284

**Authors:** Anja Mezger, Sandy Klemm, Ishminder Mann, Kara Brower, Alain Mir, Magnolia Bostick, Andrew Farmer, Polly Fordyce, Sten Linnarsson, William Greenleaf

**Affiliations:** Department of Genetics, Stanford University, Stanford, CA, USA; Department of Medical Biochemistry and Biophysics, Karolinska Institute, Stockholm, Sweden; Takara Bio USA, Mountain View, CA, USA; Department of Bioengineering, Stanford University, Stanford, CA, USA; ChEM-H Institute, Stanford University, Stanford, CA, USA; Chan Zuckerberg Biohub, San Francisco, CA, USA; Department of Applied Physics, Stanford University, Stanford, CA, USA

## Abstract

We have developed a high-throughput single-cell ATAC-seq (assay for transposition of accessible chromatin) method to measure physical access to DNA in whole cells. Our approach integrates fluorescence imaging and addressable reagent deposition across a massively parallel (5184) nano-well array, yielding a nearly 20-fold improvement in throughput (up to ~1800 cells/chip, 4-5 hour on-chip processing time) and cost (~98¢ per cell) compared to prior microfluidic implementations. We applied this method to measure regulatory variation in Peripheral Blood Mononuclear Cells (PBMCs) and show robust, *de-novo* clustering of single cells by hematopoietic cell type.

---

A central challenge of systems biology is to determine the epigenome of phenotypically distinct cellular states within complex primary tissue. Towards this goal, single-cell chromatin accessibility measurements provide an important epigenetic view of the regulatory landscape within individual cells by capturing the physical accessibility of putative functional elements across the genome^1–5^. Methods for measuring chromatin accessibility at single-cell resolution, however, are low throughput^3–5^, depth limited^6^, or require complex molecular processing to generate cellular indexing reagents^2,6^. For ultra-high throughput accessibility profiling applications, combinatorial indexing approaches^2,6^ offer significant promise, yet these methods capture fewer accessible fragments per cell than single-cell isolation technologies^1,3^ and are not amenable to integration with single-cell microscopy or other multi-omic assays that require whole, live cells. In this report, we describe a high-throughput implementation of single-cell ATAC-seq^7^ (scATAC-seq) that directly integrates fluorescence imaging and provides an extensible foundation for multi-omic epigenetic profiling in single cells.

We have implemented scATAC-seq in small volumes using a recently developed nano-scale liquid deposition system (ICELL8 Single Cell System, Takara Bio USA). This approach reduces reagent costs and achieves equal or higher per-cell fragment counts than prior state-of-the-art implementations^2,3,6^. The workflow -- illustrated in **Figure 1a** -- is comprised of five on-chip steps: (1) isolated single cells are stained with Hoechst and propidium iodide and stochastically loaded under Poisson statistics (~1 cell per well on average) across 5184 wells under active humidity and temperature control; all wells are then imaged via multi-color microscopy to identify those containing a single live cell; (2) transposition reagents are added to a selected set of wells (e.g., those containing a single live cell) and incubated at 37°C for 30 minutes; (3) the transposition reaction is quenched by incubation with EDTA; (4) MgCl_2_ is added in equimolar concentration to quench the chelating capacity of EDTA in preparation for subsequent PCR amplification; and (5) PCR reagents are added and scATAC-seq fragment libraries are amplified using barcoded primers provided in the prior two steps (see **Supplementary Table S1** for reagent loading chart). Following on-chip library construction, indexed scATAC-seq libraries are extracted from all nano-wells by centrifugation, purified, and then further amplified as necessary for sequencing (see **Methods**).

**Figure 1.**
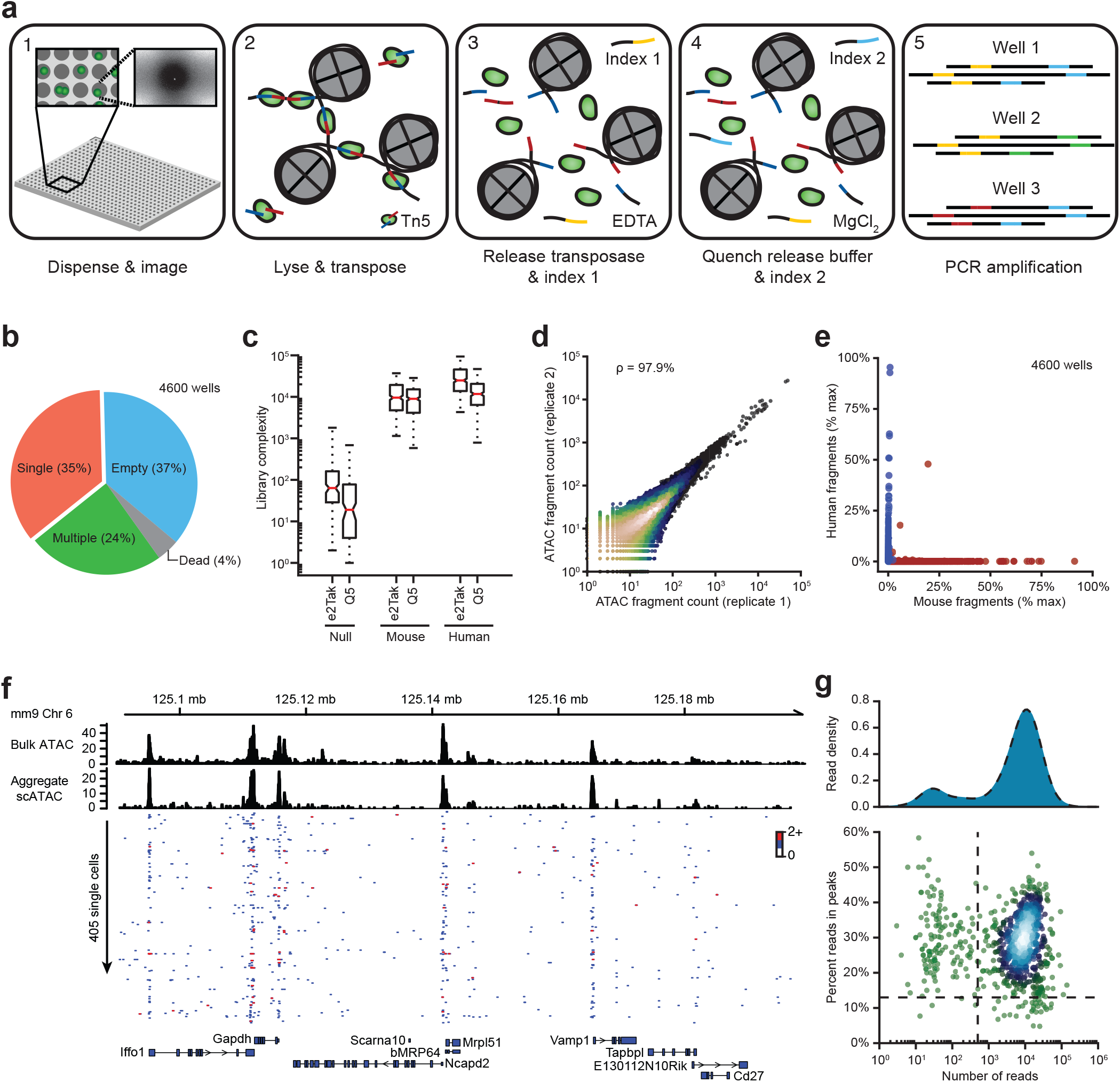
Nano-well scATAC-seq implementation on the ICELL8 platform. (**a**) Single-cell ATAC-seq workflow. (**b**) Distribution of cell counts per well measured by fluorescence microscopy (Hoechst). (**c**) scATAC-seq library complexity for null, mouse and human targeted wells using two separate polymerases (e2Tak and Q5) for well barcoding and amplification. For each sample, the box denotes the interquartile range centered at the median (red line), while the whiskers span the 5^th^ and 95^th^ percentile range. (**d**) Correlation between nano-well chips processed with either a Q5 or e2Tak polymerase across all accessible loci. (**e**) Interwell fragment contamination. (**f**) Representative population^8^ and single-cell ATAC-seq genome tracks for the Gapdh locus. (**g**) Signal-to-background (percent reads in peaks) as a function of read depth. Only cells lying in the upper right quadrant (defined by dashed lines) are retained for subsequent analysis.

As an initial test of the proposed scATAC-seq implementation we loaded samples into 5,000 wells across two nano-well ICELL8 chips. On each chip, 200 wells were loaded with PBS (designated null wells) and 1,150 wells were loaded with either mouse embryonic stem cells (mESCs) or human lymphoblastoid GM12878 cells. Imaging of Hoechst and propidium iodide fluorescence revealed the anticipated fraction of wells containing live single cells (35%, 1610 single cells), consistent with near optimal loading that maximizes the number of single-cell containing wells (**Figure 1b**). Barcoded sequencing of each of the 5,000 targeted wells revealed 14,321 (median) fragments per cell, reflecting two orders of magnitude enrichment of mappable sequences relative to null wells (**Figure 1c**). Purified nuclei from mESCs yielded marginally lower library complexities, reflecting chromatin loss during nuclear isolation (10,514 median fragments per cell). For both cell types, we observe a >10-fold enrichment for fragments proximal to transcription start sites (TSS) relative to distal regions, reflecting a high ratio of fragments captured within open rather than closed chromatin (**Supplementary Figure S1a**). Furthermore, we find a high degree of concordance (97.9%) between nano-well chips even when scATAC-seq fragments are amplified with different polymerases (**Figure 1d**). Given the high degree of deposition control on the ICELL8 platform, the inter-species crosstalk between wells was measured to be less than 0.1% (**Figure 1e**).

Aggregate single-cell profiles recapitulate population measurements broadly across the accessible genome (**Supplementary Figure S1b**) as well as specifically at individual genomic loci (**Figure 1f**). At single-cell resolution, accessibility profiles are enriched for open chromatin (**Figure 1f,g**) in both mESCs (29% reads in peaks, **Figure 1g**) and GM12878 cells (22% reads in peaks, **Supplementary Figure S1c**). Collectively, these data establish the proposed nano-well implementation as a high-throughput framework for scATAC-seq library construction.

We next asked whether scATAC-seq epigenetic profiles are sufficient to distinguish cell types within complex primary tissue. For this purpose, we performed scATAC-seq on human Peripheral Blood Mononuclear Cells (PBMCs) as well as B, T, CD4+ T, CD8+ T and monocyte cells isolated directly from whole blood (**Figure 2a**), yielding 2333 single cells passing all quality control criteria (see **Methods**). Using ChromVar, a bioinformatic approach described previously^9^, we calculated the relative accessibility of transcription factor binding motifs in individual cells and found that isolated B, T and monocyte cells robustly cluster by cell type (**Supplementary Figure S2a**). By aggregating fragments within single cells that are proximal to a transcription factor motif, this epigenetic signature captures the variation in putative transcription factor binding site accessibility across a population of cells^9^. A relatively small fraction of cells are incorrectly assigned to clusters; however, the frequency of these events as well as the random distribution of these cells within apposing clusters both suggest that isolation impurity upstream of the scATAC-seq assay is the primary source of these errors (**Supplementary Table S2**). PBMC subpopulations cocluster precisely with the isolated cell types (**Figure 2b,c**), showing highly concordant cell-type specific accessibility patterns within appropriate tSNE^10^ (t-Distributed Stochastic Neighbor Embedding) clusters (**Figure 2c**) as well as k-means clustering across highly variable transcription factor binding motif accessibility patterns (**Figure 2b**). Consistent with published gene expression data, we find that the PU.1 binding motif is differentially accessible in monocytes and B cells relative to T cells (**Figure 2c**, upper right panel)^11,12^, the C/EBPα motif is exclusively accessible in monocytes (**Figure 2c**, lower left panel)^13,14^, and RUNX1 motif accessibility is appropriately enriched in T cells – reflecting the broad regulatory role of the RUNX protein family in T lymphocytes (**Figure 2c**, lower right panel)^15^. These results are highly robust to biological (3 human blood donors) and technical variation (**Supplementary Figure S2b**). To further establish the robustness of clustering by cell type, we independently purified CD4+ and CD8+ T cells and found that these subtypes co-cluster with independently isolated T cells (**Figure 2c**, upper left panel). Collectively, these data suggest that scATAC-seq signatures are sufficient for *de-novo* clustering of PBMCs by hematopoietic cell type.

**Figure 2.**
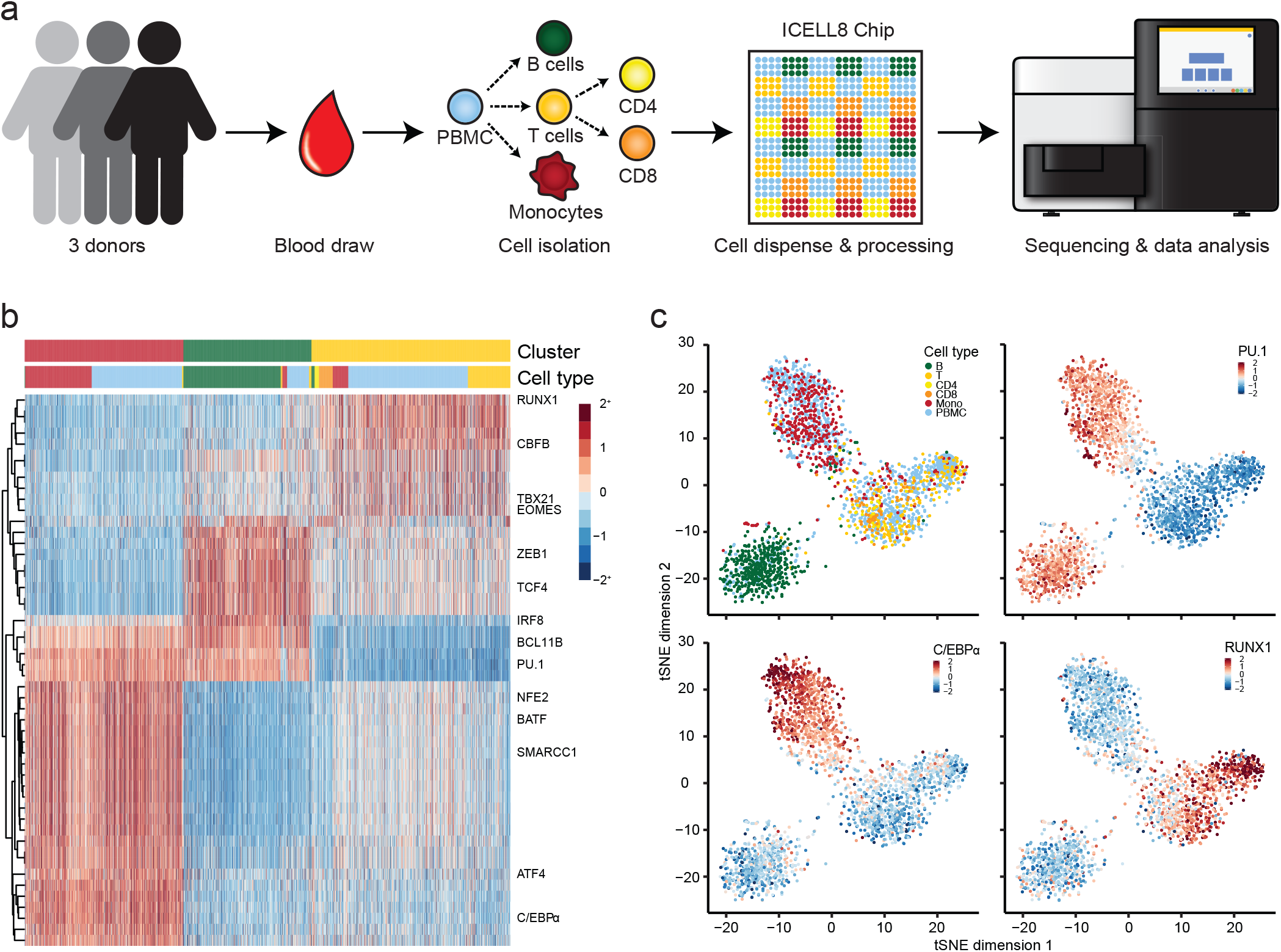
De-novo identification of hematopoietic cell types by scATAC-seq. (**a**) Human PBMC isolation and scATAC-seq workflow. (**b**) Hierarchical (TF motifs) and k-means (cells) clustering of accessibility deviation z-scores across 2333 single cells (columns) of the 50 most variable TF motifs (rows). Colors correspond to cell types defined in (**a**). (**c**) tSNE visualization of accessibility deviations at TF motifs. Cells are either colored by cell type (upper left panel) or by accessibility deviation z-score for the specified TF motif.

In this report, we have described a high-throughput implementation of scATAC-seq that dramatically reduces per-cell costs, requires only commercially available reagents, provides state-of-the-art data quality, and increases throughput nearly 20-fold over existing single-cell capture technologies. In general, nano-well single-cell sequencing approaches -- such as the method outlined here -- are highly extensible and flexible, well-suited for multi-omic analysis, and thus define an important direction for single-cell epigenetic methods development.

## Methods

Cell culture. GM12787 cells were cultured in RPMI 1640 media supplemented with L-glutamine (ThermoFisher Scientific, MA, USA, Cat. # 11875-085) and 10% FBS (ThermoFisher Scientific, Cat. # 10082147) and mESC cells were cultured in 15% FBS (HyClone GE Healthcare Life Sciences, SH30070.03E) supplemented with non-essential amino acids, L-glutamine and Leukemia Inhibitory Factor (LIF, Invitrogen, Cat. # <u>A35935</u>), respectively, at 37 °C in 5% CO2. Adherent mESCs were washed twice in PBS and detached using trypsin (Sigma, MO, USA) for 5 min. Cells were collected by centrifugation at 1000 rpm for 5 min and resuspended in their respective media. Prior to dispense, harvested cells were washed twice in media.

### Immune cell isolation from whole blood

Monocytes, T cells, CD4 T cell, CD8 T cells and B cells were isolated from whole blood using the respective EasySep Direct Human cell isolation kit (STEMCELL Technologies, MA, USA). Isolation was performed per manufacturer’s recommendation. Isolated PBMCs (AllCells, CA, USA) were thawed in RPMI and washed once in media before staining the cells as described below.

### ICELL8 workflow

Cells were stained with Hoechst and propidium iodide using the ReadyProbes Cell Viability Imaging Kit (ThermoFisher Scientific) for 20 min in media at 37 °C, then washed twice in cold 0.5x PBS. Cells were counted and dispensed into nano-wells using the ICELL8 MultiSample NanoDispenser (MSND, Takara Bio USA, CA, USA) at 25 cells/µl in 0.5x PBS, 1x Second Diluent (Takara Bio USA) and 0.4 U/µl RNase Inhibitor (New England Biolabs [NEB], MA, USA) into a blank 250 nl v-bottom deep ICELL8 chip (Takara Bio USA). Control wells containing PBS and fiducial mix (Takara Bio USA), respectively, were included in the source loading plate. All depositions were performed as 40 nl additions. Following cell deposition, chips were sealed with imaging film (Takara Bio USA) and centrifuged at 500 g for 5 min at 4 °C and imaged with a 4x objective using Hoechst and propidium iodide fluorescence. Immediately following imaging, the Tn5 transposition mix (2x TD buffer [20 % Dimethylformamide, 20 mM Tris-HCl, pH 7.6, 10 mM MgCl_2_, 100 µl Tn5 transposase [Nextera DNA Library Prep Kit, Illumina, CA, USA] per ml Tn5 transposition mix, 0.2 % Tween20, 0.2% NP40, and 0.02 % Digitonin [Promega, WI, USA]) was dispensed. Chips were then sealed, centrifuged and incubated for 30 min at 37 °C. To index the whole chip, 72 i5 and 72 i7 previously published, custom indexes^3^ were dispensed with EDTA and MgCl2, respectively. To release the bound Tn5 transposase, 60 mM EDTA and 6.25 µM custom Nextera i5 indexes were dispensed. After sealing the chip, the chip was centrifuged at 3,000 g for 3 min and incubated for 30 min at 50 °C. In order to quench the high EDTA concentration before performing PCR on-chip, 60 mM MgCl2 was dispensed together with 6.25 µM of the i7 indexes. Chips were then sealed, centrifuged, and incubated at room temperature for 5 min. Finally, a PCR mix (5x Q5 [NEB] or e2TAK [Takara Bio USA] reaction buffer, 1 mM dNTPs [Thermo Fisher Scientific], and 0.1 U/ml Q5 [NEB] or e2TAK polymerase [Takara Bio USA], respectively) was dispensed and 14 cycles of PCR were performed on-chip after sealing and centrifuging at 3,000 g as follows: 5 min at 72 °C and 30 sec at 98 °C followed by 14 cycles of 10 sec at 98 °C and 90 sec (Q5 polymerase) or 150 s (e2TAK polymerase) at 72 °C, with a final extension of 2 min at 72 °C. PCR products were extracted by centrifugation at 3,000g for 3 min using supplied extraction kits (Takara Bio USA).

### Off-chip purification and additional amplification

The collected PCR product was purified using MinElute PCR purification columns (Qiagen, Germany) following the manufacturer’s instructions. Due to the large sample volume, the PCR product was split across four MinElute columns, eluted in 10 µl volumes, and subsequently pooled. To remove free PCR primers, which would induce index-swapping during additional rounds of amplification, we performed two rounds of bead clean-up using Ampure XP beads (Beckman Coulter, CA, USA) in a 1:1.2 ratio. The beads were incubated for 8 minutes with the PCR product, washed twice in 70% ethanol, and eluted in 20 µl ultrapure water (Thermo Fisher Scientific). Further amplification was required only for the mouse and human mixing experiment; on-chip generated PBMCs libraries were directly sequenced following column and bead purifications.

The number of off-chip amplification cycles was determined by running a 20 µl qPCR reaction (2 ul PCR product, 0.5 µM oligo C [Illumina P5], 0.5 µM oligo D [Ilumina P7], 0.6 x SYBR Green I [Thermo Fisher Scientific], and 1 x NEBNext High-Fidelity 2X PCR Master Mix [NEB]): 5 min at 72°C, 30 sec at 98°C, followed by 20 cycles of 10 sec at 98°C and 90 sec at 72°C. The number of cycles that corresponded to 1/3 of the maximum fluorescence intensity was determined and the remaining 18 µl of PCR product were amplified according to that number using the same PCR conditions. The amplified PCR product was cleaned up using a Qiagen MinElute column.

### DNA sequencing

All libraries were sequenced on a NextSeq500 (Illumina) using the high output v2 kit (Illumina) in 76 x 8 x 8 x 76 cycle mode, although 36 bp x 8 x 8 x 36 bp sequencing is sufficient. On average, approximately 50K reads were sequenced per cell. Due to the nature of the sequencing libraries 30% - 40% phiX control v3 (Illumina) was spiked in and 1.5 pM were loaded onto the flow cell according to manufacturer’s instructions.

### Per cell cost estimate

The per cell library preparation cost is conservatively estimated (assuming only 1200 single cells captured per chip) at 98¢/cell: (1) Takara Bio ICELL8 chip (52¢/cell), (2) Illumina Tn5 (24¢/cell), (3) e2Tak polymerase (4¢/cell). The per cell sequencing cost at the depth used for this report is approximately 17¢/cell; other reagents contribute less than 1% additionally, yielding a total scATAC-seq cost of 98¢ per cell.

### Data analysis

Illumina sequencing reads in BCL format were demultiplexed by single-cell barcode to fastq files using bcl2fastq (Illumina) according to the manufacturer’s manual. Reads were trimmed using Cutadapt^16^ (parameters: -a Trans2_rc = CTGTCTCTTATACACATCTCCGAGCCCACGAGACA, Trans 1_rc=CTGTCTCTTATACACATCTGACGCTGCCGACGA) and aligned to either the human (hg19) or mouse (mm9) genomes using Bowtie2^17^. Mitochondrial reads were removed prior to downstream analysis. PCR duplicates were identified and removed if either the start or end position was shared with another sequencing read. Library complexity estimates were performed using Picard Tools MarkDuplicates utilities (https://broad-institute.github.io/picard/). Accessible chromatin regions (peaks) were determined using MACS2^18^ (parameters: --format BAMPE --nomodel --call-summits --nolambda --keep-dup all) for mouse embryonic stem cells (mESCs) and human lymphoblastoid (GM12878) cells. A previously published accessible peak set for hematopoiesis was used for PBMC, T and B cell analysis^1^. Single cells were selected based on imaging using the supplied ICELL8 CellSelect software (Takara Bio USA) with manufacture settings. Primary PBMCs with fewer than than 500 unique (non-mitochondrial) reads or with less than 20% (10-15% for mESCs and GM12878 cells) of mappable reads lying within peaks were eliminated from subsequent analysis. Bias-corrected deviations in accessibility near transcription factor motifs were calculated using ChromVar^9^. Hierarchical and k-means clustering as well as tSNE analyses^10^ were completed using custom R scripts.

## Acknowledgments

Anja M. is supported by the Swedish Research Council (grant 2015-06403). S.K. is supported by a T32 Ruth L. Kirschstein National Research Service Award (Institutional Training Grant in Genome Science NIH 5 T32 HG000044). This work was supported by NIH (P50HG007735 and UM1HG009442 and U19AI057266 to W.J.G.), the Rita Allen Foundation (to W.J.G.), the Baxter Foundation Faculty Scholar Grant, and the Human Frontiers Science Program grant RGY006S (to W.J.G), and the Joint Institute for Metrology in Biology. W.J.G is a Chan Zuckerberg Biohub investigator and acknowledges grants 2017-174468 and 2018-182817 from the Chan Zuckerberg Initiative.

